# Transgenesis enables mapping of segmental ganglia in the leech *Helobdella austinensis*

**DOI:** 10.1101/2024.02.14.580271

**Authors:** Dian-Han Kuo, Lidia Szczupak, David A. Weisblat, Enrique L Portiansky, Christopher J. Winchell, Jun-Ru Lee, Fu-Yu Tsai

**Affiliations:** Dept. Life Science, National Taiwan University, Taipei, Taiwan; Museum of Zoology, National Taiwan University, Taipei, Taiwan; Departamento de Fisiología, Biología Molecular y Celular, Facultad de Ciencias Exactas y Naturales, Universidad de Buenos Aires and IFIBYNE UBA-CONICET. Pabellón II, piso 2. Ciudad Universitaria. C1428EHA Buenos Aires. Argentina; Dept. of Molecular & Cell Biology, University of California, Berkeley, USA; Laboratory of Image Analysis, School of Veterinary Sciences, National University of La Plata, CONICET. Argentina; 6Max-Planck-Institut für Biophysikalische Chemie, Göttingen, Germany; 7Genetics, Genomics and Development, Cornell University, Ithaca, NY, USA

## Abstract

The analysis of how neural circuits function in individuals and change during evolution is simplified by the existence of neurons identified as homologous within and across species. Invertebrates, including leeches, have been used for these purposes in part because their nervous systems comprise a high proportion of identified neurons, but technical limitations make it challenging to assess the full extent to which assumptions of stereotypy hold true. Here, we introduce Minos plasmid-mediated transgenesis as a tool for introducing transgenes into the embryos of the leech *Helobdella austinensis* (Spiralia; Lophotrochozoa; Annelida; Clitellata; Hirudinida; Glossiphoniidae). We identified an enhancer driving pan-neuronal expression of markers, including histone2B:mCherry, which allowed us to enumerate neurons in segmental ganglia. Unexpectedly, we find that adult *Helobdella* ganglia contain fewer and more variable numbers of neurons than in previously examined leech species.

## INTRODUCTION

The stereotypy of individually identified neurons has made invertebrate species with simple nervous systems useful for understanding the operational principles of neurons and neural circuits. In such nervous systems, landmark neurons, individually identified by location, morphology (especially size) and electrophysiological properties, can be recorded and manipulated to characterize their roles in neural circuits. Further, when the stereotypy extends beyond species boundaries, homologous neurons from different species can be compared to study the evolution of neural circuits. Assessing the stereotypy of neuronal elements beyond the easily recognizable landmark neurons has been difficult due to technical limitations, however. Closing this knowledge gap will help us better understand the plasticity and robustness of these simple nervous systems.

Among the classic invertebrate models, leeches are clitellate annelids, i.e., segmented worms in the superphylum Spiralia. The leech body plan arises from 32 segmental primordia (plus additional non-segmental tissues), organized into four fused rostral/head segments (R01-R04), 21 midbody segments (M01-M21), and seven caudal/tail segments (C01-C07). The leech nervous system is also segmentally organized; each midbody segment bears a discrete ganglion that is homologous to those in other segments, and the fused head and tail segments contain ganglionic masses composed of fused segmental ganglia (Shankland and Savage, 1997).

Two families of leeches have been used as models for neurobiology and embryology, respectively, since the 19th century (Retzius, 1891; Retzius, 1898; Whitman, 1878; Whitman, 1887). Hirudinid species (e.g., *Hirudo medicinalis* and *H. verbana*) are well-suited for studying cell physiology and the neural substrates of behavior; their 32 segmental ganglia comprise identified neurons that are accessible for intracellular recording (Szczupak, 2023; Wagenaar, 2015). Leech nervous systems are also of interest for comparative analysis for two reasons. First, the leech nervous system (∼400 neurons per ganglia; ∼12,000 in CNS) is intermediate in complexity between those of more intensively studied ecdysozoan models, i.e., *Caenorhabditis elegans* (302 neurons total in the hermaphrodite) and *Drosophila melanogaster* (∼100,000 neurons in the brain alone). Second, as spiralians, leeches represent a group that has been evolving independently of the ecdysozoans for ∼600 MY; thus, comparisons between leeches and other, distantly related organisms (e.g., *Drosophila* and vertebrates) should help to illuminate aspects of neurobiology and behavior that are deeply conserved and also those that are evolutionarily plastic.

Unfortunately, the early embryos of *Hirudo* are small and have never been cultured out of the cocoon, which makes them intractable for the intracellular injection of reagents for cell lineage tracing or genetic manipulation. For such work, glossiphoniid species (e.g., *Helobdella austinensis*; Fig. 1A, A’) are useful because their embryos are easily cultured in the laboratory and because the large, identifiable blastomeres in their early embryonic stages (Fig. 1B-1D and 1F) are amenable to a variety of experimental approaches, including intracellular injection (Kuo et al., 2020; Weisblat, 2022; Weisblat and Kuo, 2009; Weisblat and Kuo, 2014). On the other hand, the small size of its adult nervous system limits physiological approaches to studying its nervous system.

**Figure 1.**
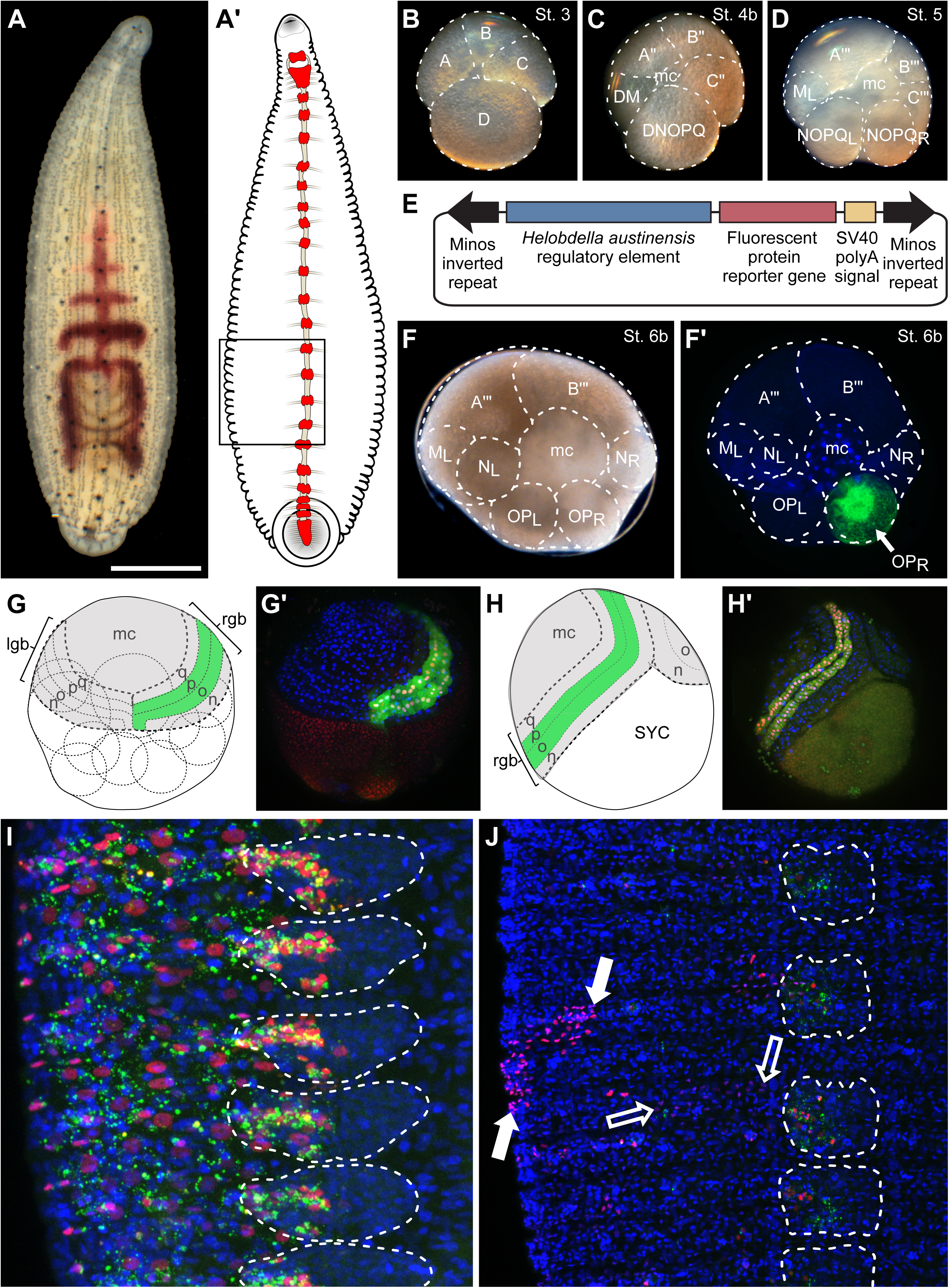
Minos-mediated transgenesis in the leech *Helobdella*. **A.** Photomicrograph of a small adult. Dorsal view, anterior up, showing dorsal eyespots at the anterior end and part of the posterior sucker at the posterior end; segmentation is evident from the repeated pigment cells in dorsal ectoderm and by articulations of the crop and intestine (visible as purple through the body wall). **A’**. Diagram highlighting the central nervous system (ventral view; ganglia of midbody segments, head and tail brains are red). **B-D.** Photomicrographs of embryos at early stages. Individual cells or cell clusters are indicated by dotted outlines. In the 4-cell stage (B), the larger, D macromere is the precursor of segmental mesoderm (including germline) and ectoderm. At fourth cleavage (C), an obliquely equatorial division of macromere D’ yields precursors of mesoderm (DM) and ectoderm (DNOPQ); in addition all four quadrants have contributed small cells to the micromere cap (mc) at the animal pole. By stage 5 (D), DM and DNOPQ have divided to establish bilateral precursors of mesoderm (only teloblast M_L_ is seen in this view) and ectoderm (proteloblasts NOPQ_L_ and NOPQ_R_), plus additional micromeres. **E.** Schematic of the Minos plasmid used for transgenesis. **F-J. Diagrams and confocal images of embryos** fixed and counterstained with Hoechst 33342 (blue) at various times after stage 6b, in which OP_R_ proteloblasts were injected with a mixture of fluoresceinated dextran amine (FDA; green) and transgenesis cocktail containing the plasmid pMi{Htr-ef1a-h2b:mCherry}(red). **F, F’.** At stage 6b (t∼0), the NOPQ proteloblasts have cleaved to form precursors contributing primarily to ventral (N teloblasts), dorsal (Q teloblasts, not visible) and lateral (OP proteloblasts) ectodermal domains. **G, G’.** By early stage 8 (t ∼48 hrs), all ten teloblasts (fine dotted outlines), lie beneath the surface, dividing asymmetrically to generate segmental founder cells in discrete columns (superficial bandlets n, o, p, q are depicted; m bandlets lie underneath) that merge at the surface into left and right germinal bands (lgb, rgb; coarse dotted outlines), then move ventrovegetally over the surface of the embryo. Germinal bands and the territory behind them is covered by a squamous epithelium (gray) derived from the micromere cap (mc). Cells in the o and p bandlets of the rgb have inherited FDA (green) and are beginning to express H2B:mCherry in their nuclei, which appear pink/white because FDA is also in nuclei at this time. **H, H’.** Obliquely ventral view of a late stage 8 embryo (t ∼96 hrs). Teloblasts and macromeres have fused to form a syncytial yolk cell (SYC) from which definitive endoderm will form. Germinal bands are coalescing anteroposteriorly along the ventral midline. Expression of H2B:mCherry is uniform within each bandlet, presumably reflecting plasmid-driven expression. In this specimen, expression is stronger in the p bandlet, presumably reflecting a non-uniform distribution of plasmid when OP divided to form O/P teloblasts. **I.** At stage 10 (t ∼9 days), showing part of six segments, corresponding to the boxed region in panel A’, the injected lineage tracer marks cells in lateral epidermis and segmental ganglia (dotted white outlines); FDA is no longer distributed continuously throughout the cytoplasm, making it harder to perceive contours of OP-derived cells. In contrast, H2B:mCherry clearly marks the nuclei of seemingly all OP-derived cells, still reflecting plasmid-driven expression. **J.** In juveniles (t ∼30 days), FDA has disappeared except for traces in ganglionic cells, which exit mitosis earlier than epidermal cells. H2B:mCherry expression is no longer uniform, reflecting the cessation of plasmid-driven expression. Instead, nuclear expression persists in subsets of ganglionic cells and patches of epidermis, indicating mosaic integration of the transgene. Contiguous patches of brighter (closed arrows) and fainter (open arrows) epidermal cells suggest differences in copy number and/or position of transgene integration. Scale bars, 1 mm in **A**, 270 μm in **B**-**D**, 225 μm in **G**-**H’**, 30 μm in **H**, and 80 μm in **J**. Animal pole (prospective dorsal) views for panels B-D, F and G. Ventral views for panels H-J.

To address those limitations and to establish groundwork for future developmental and neurophysiological studies, we have developed efficient and lineage-specific transgenesis in *Helobdella* embryos. For this purpose, we microinjected individual blastomeres of the early embryo with a cocktail of a Minos transposon-bearing plasmid and Minos transposase mRNA. Injecting germline precursors allowed us to generate lines of *Helobdella* that stably express selected transgenes. Combined with the judicious identification of a pan-neuronal enhancer, we have used Minos transgenesis to determine the neuronal content and variability in ganglia of the ventral nerve cord in *Helobdella*.

## MATERIALS AND METHODS

### Animals, embryos, microinjections

*Helobdella austinensis* embryos were obtained from a long-term laboratory colony, staged and cultured all as described elsewhere (Weisblat and Kuo, 2009). Plasmids were pressure-injected into identified blastomeres at stages 4c-7 (DM blastomeres, OP proteloblasts, or M teloblasts), using the same procedures as for microinjecting HRP and derivatized dextran lineage tracers (Weisblat and Kuo, 2009). To optimize survival, injected embryos were frequently cultured through stage 8 in HL saline containing roughly 1% commercial penicillin-streptomycin solution (Gibco).

### Screening and breeding transgenic animals

In the present experiments, injected embryos (F0 animals) were screened under a fluorescence dissecting microscope to visualize neuronal expression of fluorescent protein (FP) driven by the *Hau-elav* upstream regulatory elements at stages 9-10, when embryonic mobility was still minimal and yet neuronal differentiation was well under way. FP-expressing individuals from injected clutches were cultured further to the end of stage 11 (yolk exhaustion) and then hand-reared to reproductive maturity on a diet of small physid or planorbid snails. These F0 animals were allowed to interbreed; the resulting clutches of F1 animals comprised mainly clutches with no expression and a few clutches of mixed expression and non-expression. FP-expressing F1 animals were selected to interbreed. Since the occurrence of FP-expressing F1 animals in each clutch is typically low (0%-29%) and because *Helobdella* juveniles survive better when cultured as groups, we often pooled FP-expressing F1s to improve survival rates and preserve transgenes within the population. The resultant F2 clutches usually contained a higher proportion of FP-expressing animals; this allowed us to select and grow cohorts of FP-expressing F2s from individual clutches in isolation to establish inbreeding lines. We scored the proportion of FP-expressing animals among the F3 clutches to estimate the inheritance pattern of transgenes. F3 clutches with 100% transgene expression generally gave rise to progeny consistently expressing the FP reporter.

### Identification and characterization of *Hau-elav* genes

Four *elav* homologs were identified from the InterPro *H. robusta* proteome dataset (Accession: UP000015101) using the query IPR006548, an InterPro ID that structurally defines the Elav/Hu family members. The amino acid sequences of the four *H. robusta* elav proteins retrieved from the InterPro database were used to perform TBLASTN searches of gene models in the *H. robusta* genome database (JGI, DOE, USA) and *H. austinensis* transcriptomes to obtain their nucleic acid sequences. For each of these four *Hau-elav* genes, an ortholog was identified in *H. robusta*.

Phylogenetic analysis of Elav proteins from selected invertebrate species was performed using additional sequences retrieved from the EnsemblMetazoa database. The amino acid sequences were aligned using the Muscle algorithm (Edgar, 2022). Maximum-Likelihood tree search was performed using IQ-TREE (Nguyen et al., 2015); the best-fit substitution model was identified by ModelFinder (Kalyaanamoorthy et al., 2017) and automatically implemented during the tree search sessions using the optional argument -m MFP in IQ-TREE; 1,000X approximated bootstrapping was performed using the ultrafast bootstrap algorithm (Hoang et al., 2018). The plasmid templates for synthesizing antisense riboprobes of the *H. austinensis elav* genes (*Hau-elav1*, *Hau-elav2*, *Hau-elav3*, and *Hau-elav4*) were produced by PCR cloning of cDNA fragments. The primer pairs for amplifying these cDNA fragments are shown in Table S1. Probe synthesis and whole-mount *in situ* hybridization were performed as previously described (Weisblat and Kuo, 2009).

### Transgenic reporter constructs

Genomic organization for both *Hau-elav4* and *Hau-cif7* was reconstructed by mapping the *H. austinensis* genome assembly with the EST-supported *H. robusta* gene models and *H. austinensis* transcripts recovered from the transcriptomes. The genomic organization of ortholog pairs is conserved between *H. robusta* and *H. austinensis*. The upstream genomic fragments of *Hau-elav4* and *Hau-cif7* were PCR amplified using primers listed in Table S1 and cloned into pJET vector. The resulting plasmids contain a 3,552 bp fragment corresponding to -3,549 to +3 bp relative to the *Hau-elav4* translation initiation site and a 4,279 bp fragment corresponding to -4,255 to +24 bp relative to the *Hau-cif7* translation initiation site, respectively. These two plasmids served as the starting materials for subsequent reporter constructs.

The base of the reporter constructs was modified from the pMi{3XP3-dsRed} plasmid (Pavlopoulos and Averof, 2005). Briefly, the 3XP3-dsRed insert was excised by restriction digestion with *Bam*HI, followed by self-ligation. The resulting pMinos plasmid contains several unique restriction sites that can be used to insert DNA fragments between the left and right *Minos* elements.

A DNA fragment containing the SV40 polyadenylation signal was excised from pCS107 vector by double digestion with *Xho*I and *Asc*I and inserted into pBSMNEF1P (Gline et al., 2009), resulting in pEF-SV40. Next, the DNA fragment containing the 2,270 bp *Helobdella triserialis* EF1α upstream sequence and the SV40 element was excised from pEF-SV40 by double digestion with *Bam*HI and *Sac*II and then inserted into pMinos plasmid to yield pMi{Htr-ef1a-sv40}. *Cla*I and *Xho*I sites were introduced to the 5’- and 3’-ends of nGFP, H2B:GFP, and H2B:mCherry coding sequences by PCR amplification from pCS2+nGFP (Zhang and Weisblat, 2005), pCS107 H2B:GFP (Gline et al., 2009), and pCS107 H2B:mCherry, respectively. The restriction digested amplicons were then inserted between *Cla*I and *Xho*I sites of pMi{Htr-ef1a-sv40} to generate pMi{Htr-ef1a-ngfp} and pMi{Htr-ef1a-h2b:gfp}. pC107 H2B:mCherry was created by replacing eGFP coding in pCS107 H2B:eGFP with the mCherry coding sequence. The primer pairs for amplifying DNA fragments encoding nGFP, H2B:GFP, and H2B:mCherry are listed in Table S2.

The tissue-specific reporter constructs (pMi{Hau-elav4-ngfp}, pMi{Hau-elav4-h2b:mcherry}, and pMi{Hau-cif7-ngfp}) were produced using NEBuilder HiFi DNA Assembly kit (NEB). The pMi base plasmid was linearized by double digestion with SpeI and SacII. The upstream DNA fragments were PCR amplified from their respective plasmid templates. The DNA fragments encoding the nGFP-SV40 and H2B:mCherry-SV40 reporters were amplified from the pCS2+ nGFP and pCS107 H2B:mCherry. The combinations of primer pairs and plasmid templates for generating DNA fragments used in the DNA assembly reactions are listed in Table S3. The resulting pMi{Hau-elav4-ngfp} and pMi{Hau-elav4-h2b:mcherry} plasmids contain a 2,587 bp DNA fragment upstream to the start codon of *Hau-elav4*, followed by an nGFP-SV40 or H2B:mCherry-SV40 reporter, respectively. The pMi{Hau-cif7-ngfp} plasmids contain a 4,254 bp fragment upstream to *Hau-cif7*, followed by an nGFP-SV40.

### Analysis of transgene copy number

Genomic DNA of transgenic or wild-type leeches was individually extracted using ZR Genomic DNA Tissue Microprep Kit (ZymoResearch). Following quantification, genomic DNA was restriction digested with *Hind*III (New England Biolabs) and circularized by self-ligation. The cloned *Hau-elav4* upstream region contains a single *Hind*III site near the 5’ end, and the frequency of *Hind*III sites in the *Helobdella* genome was estimated to be one per 2,916.4 bp on average. The concentrations of genomic DNA (0.1 ng/μl) and T4 DNA ligase (0.04 U/μl) and the reaction temperature (16°C) were kept low to suppress the formation of heteroduplex DNA. After ligation, 1 ng of self-circularized genomic DNA was used as PCR template and amplified with a pair of inversely oriented primers.

The forward primer annealed to the SV40 polyadenylation signal region, and the reverse primer annealed to a region immediately downstream of the *Hind*III site in the cloned 2,587 bp *Hau-elav4* upstream DNA fragment. The sequences of this primer pair are given in Table S4. This amplicon is designed to capture endogenous genomic DNA fragments adjacent to the Minos right arm at the insertion site. Amplicons were analyzed by agarose gel electrophoresis to determine the number of insertions in each individual. Selected amplicons were sequenced to confirm genomic DNA integration and to determine the genomic location of the insertion sites.

### Dissections, histology and imaging

For the initial evaluation of the *Hau-cif7* and *Hau-elav4* reporters, the embryos injected with the transgenic cocktail were raised to stage 10 and stage 11 respectively, and fixed at 4°C overnight in a fixative containing 4% formaldehyde and 0.5X PBS. The fixed specimens were counterstained with Hoechst DNA stain. For stage 10 ganglia, the germinal plate was dissected off the yolk, flattened, and mounted in 80% glycerol. Images of stage 11 ganglia were acquired from whole-mount specimens cleared with the DeepClear protocol (Pende et al., 2020).

For high-resolution analysis of segmental ganglia, three (3) animals were dissected along the dorsal and ventral midlines, fixed in 4% formaldehyde overnight, rinsed in PBS, and cleared in 40, 60, and 80% glycerol. Finally, they were mounted in 80% glycerol for imaging. Ganglia M2 – M18 were imaged using a 20X objective on a Zeiss LSM 900 confocal microscope, ventral side up; mCherry fluorescence was excited by a 561 nm laser. Whole ganglia images were obtained using a Z-spacing of 2 µm. Individual packets were captured as 8-bit grayscale images (resolution 2048 X 2048), using a digital zoom between 2.5X and 3.8X with a Z-spacing of 0.63 µm. Data were acquired as CZI files and transformed to TIF stacks for analysis.

### Image analysis

To optimize resolution we imaged ganglia on a packet-by-packet basis. Confocal images were processed using ImagePro Plus (V6.3, Media Cybernetics, MD). Stack images were used for generating 2D or 3D images. 2D images were used to measure the area (area of the z projected nuclei), mean diameter, roundness ((perimeter^2^/(4π*area)); roundness=1 is circular), perimeter, mean intensity (average intensity of all the pixels in the object) of the fluorescent nuclei and the integrated optical density/intensity (IOD) which reports the average intensity/density of each object (nuclei) multiplied by the area of that object (Portiansky, 2018). For this purpose, the Extended Depth of Field (EDF, also known as Extended Focus Image (EFI; Olympus cellSens) or Z-projection in MIT Image (FIJI)) option was applied using the Max intensity focus analysis option. Nuclei in calibrated images were manually circumscribed and simultaneously counted and measured. To test the accuracy of the counting procedure, the neurons identified in selected packets were checked independently and found to be identical. Raw data are provided in Dataset 1.

3D images were used either as a model to identify nuclei in 2D projected imagenes as well as to measure volume parameters of some packets. 180-degree rotated rendered 3D images were used to detect nuclei hidden behind large neurons on the dorsal side of the ganglion (Movie 1). Volume parameters (volume, surface area, diameter, sphericity) were calculated using the 3D Constructor plug-in algorithm of the image analysis software. Data was exported to a spreadsheet for statistical analysis.

## RESULTS

### The transposable element Minos is an efficient tool for transgenesis in the leech *Helobdella*

A previous application of transgene-based lineage tracers in *Helobdella* used a plasmid in which a pair of I-*Sce*1 recognition sequences flanked a transgene cassette consisting of a 2,270 bp genomic DNA fragment upstream to the coding sequence of *Helobdella triserialis* elongation factor-1-alpha (*Htr-ef1a*) and a fluorescent protein (FP) reporter (Gline et al., 2009). Injecting this plasmid alone resulted in transient ubiquitous reporter expression with low mosaicism among the progeny of the injected cell in F0 embryos - making it useful for lineage tracing in the leech embryo, but coinjecting this plasmid with the transposase (I-*Sce*I meganuclease) yielded no stable integration.

As an alternative to the meganuclease approach, constructs based on the Minos transposon have been used to insert transgenes into the host genome in various arthropod and chordate species (Pavlopoulos et al., 2007). To see if the Minos system is useful in *Helobdella austinensis* - a spiralian species that is phylogenetically distant to arthropods and chordates, we generated Minos-based transgene constructs (pMi{ef1a-ngfp} and pMi{ef1a-h2b:mcherry}) by transferring the transgene cassette from the previously made *I-Sce1* plasmids into a plasmid containing the *Minos* transposable elements (Fig. 1E). We then injected early blastomeres with a transgenesis cocktail of the plasmid and mRNA encoding Minos transposase mixed with a fluoresceinated dextran tracer (Fig. 1F’) and monitored the transgene reporter expression in the injected F0 individuals and their descendants. As expected, we observed uniform expression of the transgene at early times following injection, reflecting plasmid-driven expression (Fig. 1G-1I; Gline et al. 2009). Expression at later stages became mosaic, as unintegrated plasmids were degraded, while expression from integrated transgene persisted (Fig. 1J).

### Breeding F0 individuals to generate transgenic lines

To achieve germline transmission of the transgene, we injected the transgenesis cocktail into embryonic germline precursor cells, i.e., M teloblasts or their precursors in the D quadrant lineage (Cho et al., 2014; Kang et al., 2002). In one experiment, the D’ blastomere was injected in 8-cell embryos (stage 4a). Of 42 injected embryos, 27 (64%) developed normally to adulthood, 22 of which showed readily visible fluorescence. Most of the expected progeny cells of the injected blastomere expressed FP; mosaicism was evidenced mainly by differences in the level of expression, especially within the epidermis, segmental ganglia and testes sacs (Fig. 2A, 2B). In some cases, mosaicism allowed us to infer the timing of integration events relative to the stereotyped cell division patterns of the injected lineage. For example, Fig. 2C shows a young F0 adult grown from an embryo in which the transgenesis cocktail was injected into mesodermal precursor DM shortly before its equal division to form the bilateral pair of M teloblasts – evidently, the transgene was incorporated into one M teloblast but not the other, resulting in unilateral mesodermal expression of the transgene. To establish a stable transgenic line, the transgene-expressing F0 individuals selected from the experiment described above were grown to adulthood and allowed to interbreed. 20-40% of the animals in the F1 generation exhibited seemingly ubiquitous transgene expression (Fig. 2D, 2D’). However, it is noteworthy that reporter expression was first observed only in micromeres during stage 5 (not shown) and only became ubiquitous in later developmental stages in the stable lines, suggesting that the *Htr-ef1a* element is not sufficient to drive maternal and early embryonic expression of the reporter.

**Figure 2.**
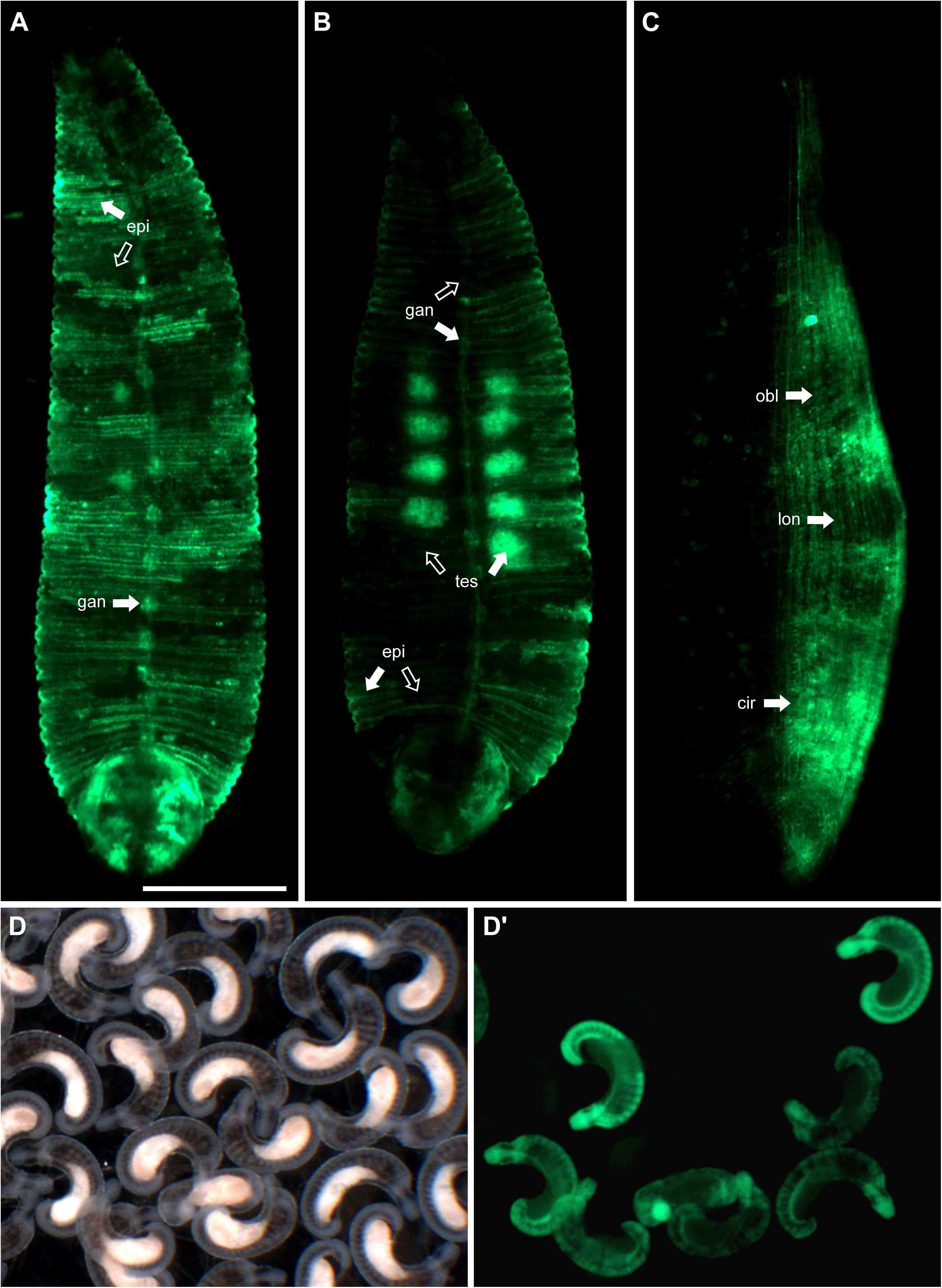
Inheritance of transgenes inserted into germline lineages. **A-B.** Fluorescence images of two F0 adults raised from embryos in which an ef1a-ngfp transgenesis cocktail was injected into macromere D’, the precursor of segmental mesoderm and ectoderm at the 8-cell stage. Mosaicism of nGFP is indicated by open arrows (little or no expression) and closed arrows (high levels of expression) in patches of epidermis (epi), segmental ganglia (gan) and testes (tes). **C.** A similar F0 adult raised from an embryo in which the transgenesis cocktail was injected into mesodermal precursor DM; DM gives rise to both M teloblasts, but transgene is evident only in circular (cir), oblique (obl) and longitudinal (lon) muscles on one side. **D and D’.** Brightfield (D) and fluorescence (D’) images of stage 10 embryos arising from interbreeding cohorts of transgenic animals such as those in (A-C). Only a fraction of the F1 cohort expresses the transgene. No mosaicism is observed but expression levels vary among individuals. Scale bar, 1 mm in A-C, 900 μm in D and D’. A-C are ventral views, with anterior up.

Initiating transgenic lines by breeding cohorts of independently injected F0 individuals means that the resulting lines should be heterogeneous with respect to the number of integrated transgenes and their chromosomal locations. Landing the transgene in different locations is expected to result in subtle differences in expression, for example from differences in the local enhancer landscape. This assumption is supported by observed differences among individuals of a breeding line bearing the ef1a-h2b:mcherry transgene, in terms of the overall brightness, and in the relative brightness of various cell types even among size-matched individuals (Fig. S1).

### Neural-specific enhancers give cell type-specific expression

Having demonstrated the feasibility of generating transgenic *Helobdella* lines using the *ef1a* enhancer, we sought to identify enhancer elements for genes expressed pan-neuronally, to implement transgenesis approaches for studying the nervous system of *Helobdella*. A previous analysis of nine cytoplasmic intermediate filament (*cif*) genes in *Helobella* revealed that *Hau-cif7* is strongly and specifically expressed in the developing nervous system (Kuo and Weisblat, 2011). In the present work, a 4,279-bp fragment upstream to the *Hau-cif7* translation start site was amplified by PCR and then cloned into the Minos transgenesis vector, yielding pMi{Hau-cif7-ngfp} (Fig. S2A). To determine the cell-type specificity of reporter expression, we injected an OP proteloblast with the transgenesis cocktail and found that the *Hau-cif7* enhancer drove transgene expression in only 3-5 cells in each hemiganglion (Fig. S2B). The number of O- and P-derived neurons in a hemiganglion is much greater than the observed five (Kramer and Weisblat, 1985; Weisblat et al., 1984), implying that the cloned *Hau-cif7* upstream region does not promote pan-neuronal expression.

Furthermore, in stage 11 embryos, the GFP-positive cells are localized on the surface of ganglionic glial packets, whereas the interior of glial packets, where the ganglionic neurons are expected, was devoid of GFP-positive cells (Fig. S2C, C’). Therefore, based on their number and distribution, these GFP-positive cells are likely ganglionic packet glial cells rather than neurons. Thus, this *Hau-cif7* regulatory element does not satisfy our requirements for this work.

The mRNA-binding protein Elav is required for neuronal differentiation in *Drosophila* and *elav* homologs have proven to be broadly conserved markers for post-mitotic neurons in various metazoan species (Colombrita et al., 2013; Pascale et al., 2008). The *Helobdella* genome contains four *elav* paralogs (Fig. 3A). Expression analysis by in situ hybridization indicates that, despite minor differences in expression patterns (Fig. 3B-E), three of the four *Hau-elav* genes are expressed in segmental and supraesophageal ganglia of developing embryos. Orthologs of these three *elav* genes are also all expressed by neurons in the adult ganglia of *Hirudo verbana* (Heath-Heckman et al., 2021). *Hau-elav4* was selected for enhancer cloning on the basis of its compact genome organization. A 2,588 bp fragment upstream to *Hau-elav4* (Fig. S2D) was amplified by PCR and cloned into the pMinos transgenesis vectors, yielding pMi{Hau-elav4-ngfp}. Injection of pMi{Hau-elav4-ngfp} transgenesis cocktail into OP proteloblast gave robust expression in previously identified neuronal elements and not in other cell types; furthermore, faintly labeled neurites were visible in advanced developmental stages (Fig. S2E, F). Thus, the cloned upstream element of *Hau-elav4* contains an enhancer capable of driving gene expression in neurons specifically.

**Figure 3.**
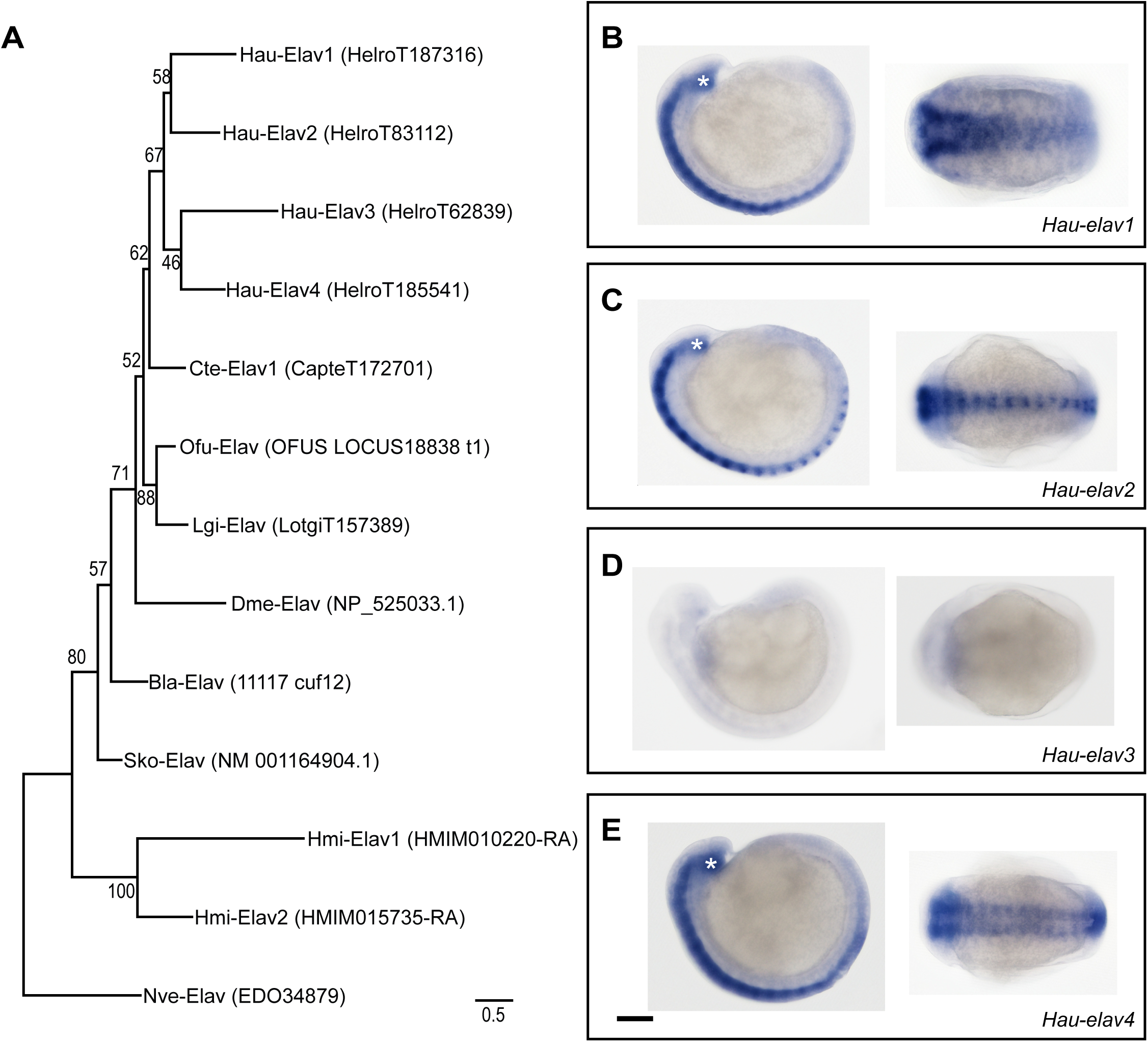
Members of elav family in *Helobdella austinensis*. **A.** Maximum-Likelihood tree of Elav proteins from selected invertebrate species. The gene model name in the EnsemblMetazoa databases for each protein is given in parentheses. Species names abbreviated as: Bla: *Branchiostoma lanceolatum* (amphioxus); Cte: *Capitella teleta* (sedantary polychaete); Dme: *Drosophila melanogaster* (fruit fly); Hau: *Helobdella austinensis* (leech); Hmi: *Hofstenia miamia* (acoel); Lgi: *Lottia gigantea* (limpet); Nve: *Nematostella vectensis* (sea anemone); Ofu: *Owenia fusiformis* (basal polychaete); Sko: *Saccoglossus kowalevskii* (acorn worm). **B-E.** Wholemount in situ hybridization revealed the expression patterns of *Hau-elav1* (B), *Hau-elav2* (C), *Hau-elav3* (D), and *Hau-elav4* (E) in stage 10 embryos. The left side of each panel shows the lateral view with dorsal to the top and anterior to the left; the right side shows the ventral view with anterior to the left. Asterisks mark the supraesophageal ganglia. Scale bar, 80 μm.

To determine whether this upstream DNA fragment of *Hau-elav4* drives pan-neuronal reporter expression, we examined the transgenic F1 progeny of F0 parents injected as embryos with a transgenesis cocktail containing pMi{Hau-elav4-ngfp} into a germline precursor cell, the M teloblast. In the successfully transgenic F1 individuals, we detected transgene expression in previously identified neuronal elements including segmental ganglia of the ventral nerve cord, the supraesophageal ganglia, and in presumptive peripheral sensory neurons of the prostomial lip and the body wall sensilli (Fig. 4A, B). Therefore, the *Hau-elav4*-driven reporter expression is likely pan-neuronal.

**Figure 4.**
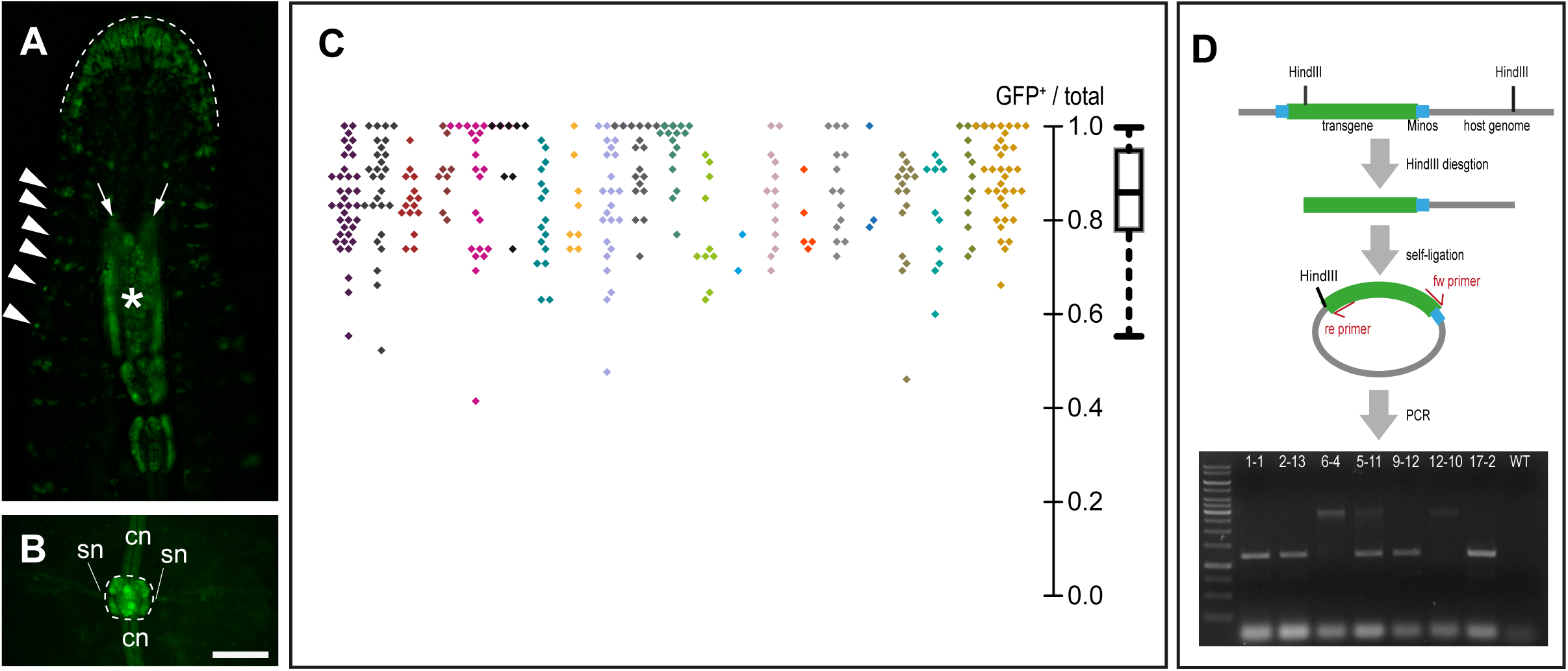
Establishment and characterization of Hau-elav4-driven transgenics. **A and B.** Fluorescence images of a transgene-positive F1 descendant of F0 animals in which an *Hau-elav4-ngfp* transgene was inserted into the embryonic germline by microinjection of M teloblasts. Ventral views, anterior is up. In the head of the animal (A), transgene expression is evident in presumptive sensory neurons of the anterior lip (dashed line), in segmentally iterated sensilli (arrowheads), and in the sub- and supraesophageal ganglia (arrows and asterisk, respectively) of the anterior ganglionic mass. In midbody segments (B) expression is high (bright signal) in neuronal cell bodies of segmental ganglia (dashed line) and is visible more faintly within connective (cn) and segmental nerves (sn), due to the imperfect nuclear localization of nGFP. **C.** Distribution of GFP-positive rates among F3 cohorts in inbreeding transgenic lines. The beeswarm plot on the left shows the ratios of GFP-positive individuals among 327 F3 cohorts drawn from 21 different interbreeding F2 cohorts, all of which arose from a single interbreeding F1 cohort. F3 cohorts derived from the same inbreeding F2 cohort are labeled with the same color. The boxplot on the right summarizes the distribution of GFP-positive rates among all scored F3 cohorts. **D.** Genotyping of selected inbreeding transgenic lines. The schematic summarizes the procedure of inverse PCR genotyping seven individuals, each from a different F3 cohort that was 100% GFP+. The image of an example gel analysis of PCR products is shown in the lower part of the panel. Scale bar, 150 μm.

To establish transgenic lines, we next crossed the transgene-expressing F1 individuals among themselves, as described above. It has been previously reported that the Minos system regularly results in multiple insertions in the host genome (Pavlopoulos and Averof, 2005). Assuming that many chromosomes would have undergone one or more transgene insertions, we anticipated that F2 cohorts derived from transgenic F1 parents would be nearly 100% positive for transgene expression. Contrary to this expectation, however, none of the F2 cohorts had such high frequencies of transgene expression, indicating that the majority of F1 individuals may have only one or two insertions per genome and were heterozygous at the insertion loci. If so, F2 cohorts derived from sibling heterozygous F1 parents should contain some homozygotes. To test this prediction, we crossed sibling F2 leeches that were positive for transgene expression and scored the rates of transgene expression in the resulting F3 cohorts. 14.68% (48/327) of F3 cohorts were found to be 100% positive, confirming that some of the F2s were indeed homozygous; the rates of transgene expression among F3 cohorts generally fell between 0.6 and 1.0, which is also consistent with the predicted inheritance pattern of 1∼2 transgene insertion loci (Fig. 4C). Overall, the expression of transgenes in our transgenic lines approximates the Mendelian ratio of a single-locus dominant allele. To verify the number of inserted transgene loci directly, we performed inverse PCR genotyping on transgenic animals taken from selected true-breeding lines. One out of the seven lines examined contained two loci, and the remaining six lines contained only a single locus (Fig. 4D). Therefore, despite having observed high success rates of transgene expression following injection of a transgenesis cocktail, which suggests a high *efficiency* of insertion, Minos transposon induced an unexpectedly small *number* of transgene insertions in the *Helobdella* genome. The factor(s) limiting the number of transgene insertions is currently unknown, but the ease of obtaining homozygotic single-locus transgenic animals is an unexpected advantage of using Minos-mediated transgenesis to study *Helobdella* biology.

### The *Hau-elav4-h2b:mCherry* transgene facilitates the analysis of neuron size and number in segmental ganglia

As described in the Introduction, each midbody segment contains a distinct segmental ganglion, numbered M01-M21 as are the segments themselves (Fig. 1A’). Within each ganglion, neurons are monopolar, with cell bodies distributed among six anatomically distinct packets: two packets make up an Anterior and Posterior pair straddling the Ventral Midline and are designated AVM and PVM respectively; four more packets constitute Left/Right pairs of Anterior and Posterior Lateral packets and are designated LAL, RAL, LPL and RPL, respectively (Fig. 5A). Within each packet, nerve cell bodies and their proximal neurites are enveloped by processes of a giant packet glia cell (Coggeshall and Fawcett, 1964). Distally, the neurites extend into the central neuropil, where they are enveloped by two other giant glia (Coggeshall and Fawcett, 1964). In addition to the giant packet and neuropil glia, ganglia also contain numerous other non-neuronal cells, including muscle cells, sheath cells, and microglia (Coggeshall and Fawcett, 1964; McGlade-McCulloh et al., 1989). Prominent longitudinal axon tracts traverse the dorsal aspect of each ganglion, continuous with the interganglionic connective nerves (Fig. 5A). Sensory and motor neurons reach the periphery through left and right segmental nerves.

**Figure 5.**
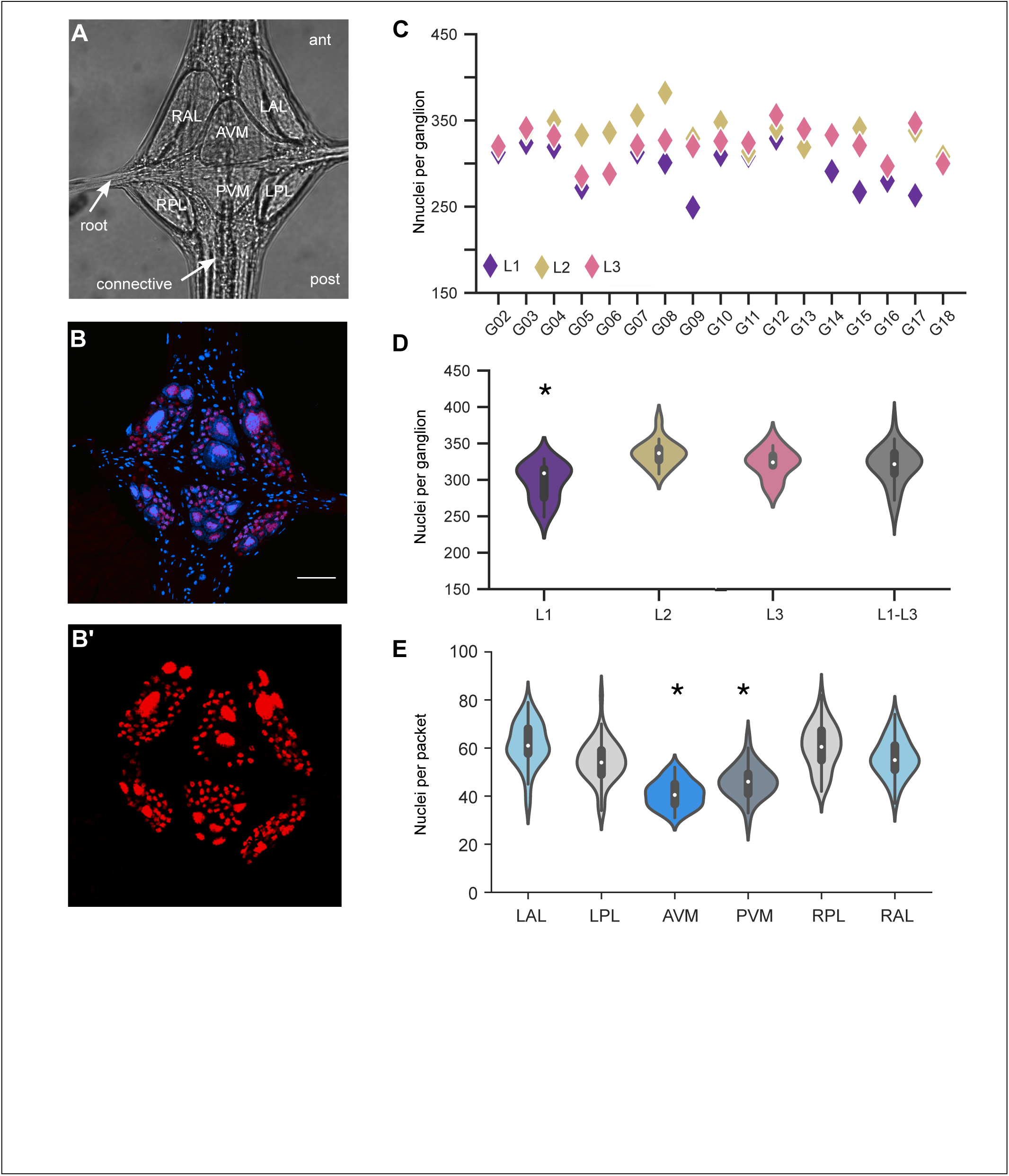
Quantification of neuronal nuclei in segmental ganglia of three leeches. **A.** Representative transillumination image of a *Helobdella* ganglion (ventral view), showing the six glial packets (RAL, RPL, AVM, PVM, LAL, PAL). Root and connective nerves are indicated. **B.** Maximum intensity projections of z-stacks of a ganglion from a Hau-elav4-h2b:mcherry transgenic leech with (**B**) and without (**B’**) DAPI staining. **C.** The count of nuclei in three leeches (L1-L3) across ganglia G02 to G18. No significant difference in the number of nuclei was found among ganglia. **D.** Violin plot showing the number of nuclei per ganglion for all the ganglia analyzed, per animal (n = 15, 16 and 17 for L1, L2 and L3, respectively) and for the three animals combined (n = 48). **E.** Distribution of nuclei across the different packets of all the ganglia analyzed (* indicates that AVM and PVM differ significantly from LAL, LPL, RPL and RAL (p < 0.000001 Kruskal Wallis, p < 0.001, post hoc Bonferroni).

In leech ganglia, neurons are identified in part by their size and location within the ganglia, and within specific packets. For example, a pair of large serotonergic neurons, known as Retzius (Rz) cells, are readily detected by their serotonin content throughout clitellates, i.e., oligochaetes and leeches (Koza et al., 2006). To assess the neuronal inventory of segmental ganglia in *Helobdella austinensis*, we determined the absolute numbers and sizes of nuclei in mid-body ganglia from three randomly selected individuals from a *Hau-elav4-h2b:mcherry* transgenic line. We used this marker for the analysis because, within ganglia, the *elav* enhancer is expressed in neurons as described above (Figures 3, 4B and 4B’). Moreover, the histone2B:mCherry fusion protein is tightly localized to chromatin of neuronal nuclei (Gline et al., 2009), making it easier to distinguish closely apposed cells (Fig. 5B’).

Based on the evidence of neural determinacy for the leech nervous systems (Muller et al., 1981), our starting hypothesis was that *Helobdella* ganglia and packets would comprise nearly invariant numbers of neurons with stereotyped size and spatial distribution, subject to possible segment-specific differences.

For each ganglion, we counted mCherry-positive neuronal nuclei on a packet by packet basis (see Materials and Methods for details). Our aim was to count neuronal nuclei in the 17 midbody segmental ganglia M02 through M18 for each of the three individuals (L1, L2, L3). Limitations imposed by the dissection protocol made it difficult to access segments anterior to M02 and posterior to M18. Of these 51 target ganglia, three ganglia could not be imaged (either at all or in part) due to obstructions arising during dissection and mounting. Thus, our final data set consisted of 48 complete ganglia (288 packets).

Our data reveal that the neuronal content of *Helobdella* ganglia is more variable than suggested by the initial hypothesis. First, visual inspection of the ganglia from any one animal suggested both similarities and differences in the distribution of neurons among different packets (Fig. 6). Moreover, comparing among the three specimens revealed that one of them had consistently lower neuron counts for most ganglia (Fig. 5C and 5D; Dataset 1), giving rise to a statistically significant difference among the three specimens (Kruskal Wallis test, p = 3.7e-05). Pairwise comparisons among the three specimens (Dunn’s test) showed that L1 differed from L2 (p = 2.4e-05) and from L3 (p = 0.01), while L2 and L3 were not significantly different from each other (p = 0.3).

**Figure 6.**
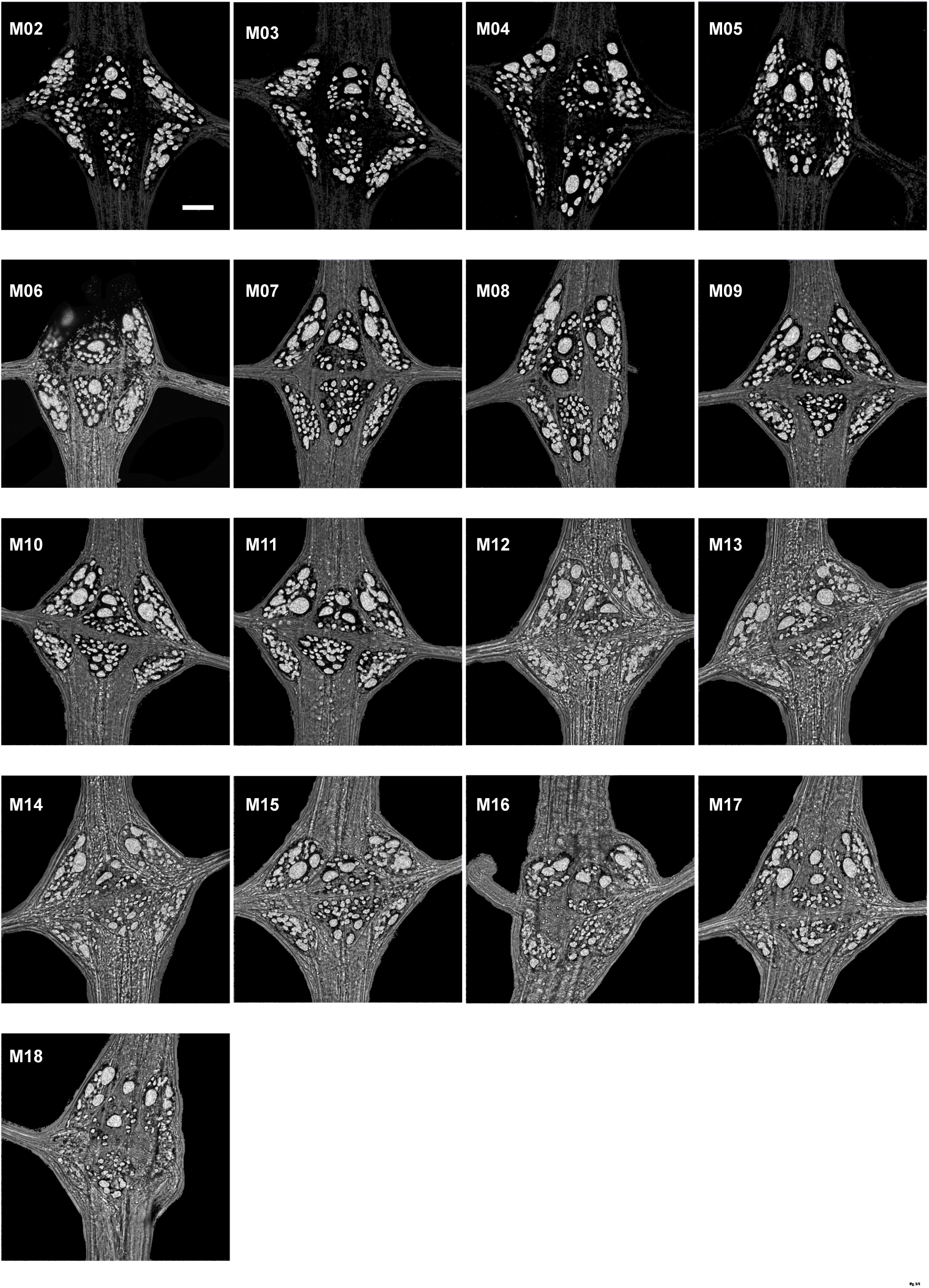
Distribution of neuronal nuclei in midbody ganglia M02 through M18 from a single transgenic individual (L3). Maximum intensity projections of z-stacks. Scale bar, 30 µm.

The total number of nuclei counted in the different ganglia varied considerably both within and among the specimens examined (Fig. 5C and 5D). This variability applied whether comparing positionally homologous segments in the three specimens, comparing serially homologous segments in each individual, or even comparing ganglia of reproductive (M5 and M6) and other segments. In the face of this unexpectedly high variability, no significant differences were observed among the neuronal complements of the 17 midbody ganglia examined here (Kruskal Wallis test, p = 0.4; Fig. 5C). Overall, the 48 ganglia span a range from 249 to 382 nuclei (319 ± 26, average ± SD) with a coefficient of variation of 0.08 (Fig. 5C and 5D).

The distribution of neurons among the six packets was also variable (Fig. 5E). However, despite the fact that the neuron counts for each of the six packets varied by as much as two-fold, we observed that the midline packets (AVM and PVM) contained significantly fewer numbers of nuclei than the lateral packets (LAL, LPL, RAL and RPL; Fig. 5E). The mean coefficient of variation for the number of neurons in the different packets is around 0.15 (0.15 ± 0.01). Thus, the distribution of neurons per packet varies more than the distribution of neurons per ganglion. This supports the possibility that neuronal cell bodies arise in somewhat predictable, but not rigidly fixed, positions during development, and that neuronal cell bodies lying between two glia end up being distributed stochastically among their respective packets during development.

Consistent with this, we observed that while AVM contains two large, presumptive Rz neurons in most of the ganglia in L3 (Fig. 6), the two large cells are distributed between the AVM and PVM packets in two ganglia (M04 and M06), and in ganglion M11, the AVM packet contains three large cells (M11). These differences in the location and number of presumptive Rz cells were not conserved in the other animals.

### Spatial distribution of large neurons shows largely conserved patterns

As described previously, another feature used to identify individual neurons in the leech ganglia is a pronounced heterogeneity in cell body size, presumably reflecting differences in the cytoplasmic volume required to meet the needs of cells with differing biosynthetic/metabolic activities, and possibly driven by differences in ploidy among post-mitotic neurons (Lasek and Dower, 1971; Yamagishi et al., 2011). For simplicity, we used the area of the 2D projection of the labeled nuclei as a proxy for volume, after first testing the validity of the correlation between projected area and measured volume in a single ganglion (Fig. S3; see Materials and Methods for details).

The projected area of the nuclei we counted varied over a >150-fold range (Fig. S4, Dataset 1). Because neurons with large somata have been identified anatomically and functionally in other species (chiefly *Hirudo* and other hirudinids, but also in *Haementeria*, a glossiphoniid; Kramer and Goldman, 1981; Kramer and Weisblat, 1985), we studied the spatial distribution of different sizes of neurons within *Helobdella* ganglion, as candidates for future identification. To correct for systematic differences among animals, the raw size data for each animal was normalized to the mean area of the largest 1% of nuclei in that animal (Fig. S3). We then defined discrete four size classes (small, medium, large and largest), based roughly on breaks in the size distribution among different packets (Fig. 7A and S3).

**Figure 7.**
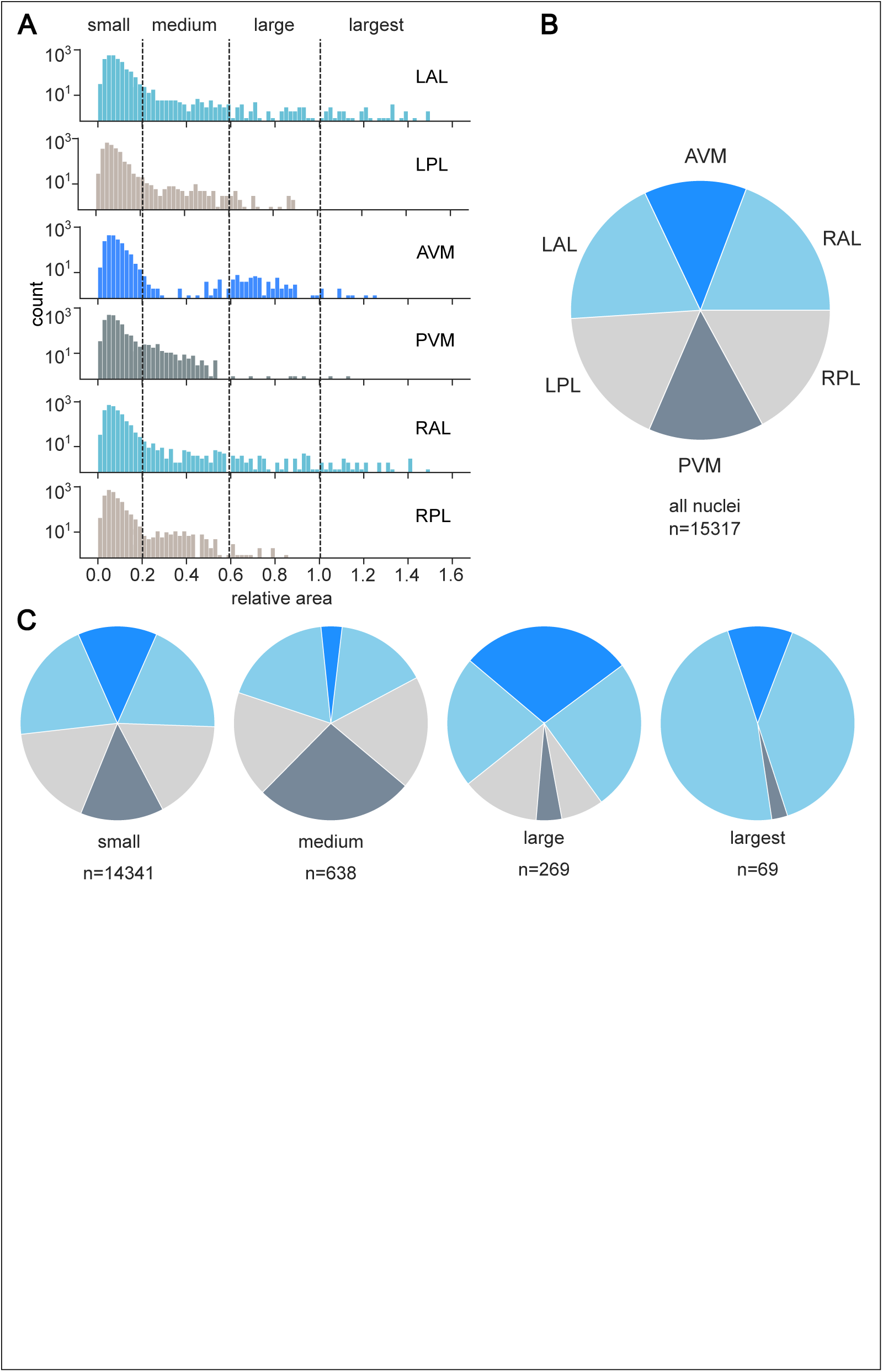
Analysis of nuclear size distribution. **A.** Histogram of relative nuclei size (measured as the 2D projection area) among packets (n = 48 ganglia). The absolute sizes of the 15,317 2D projections ranged from 3 to 537 µm^2^ in the histologically processed specimens–these raw data correspond to the nuclear chromatin of fixed and dehydrated tissue and thus are smaller than the nuclei of live neurons. The vertical lines indicate the divisions used to classify nuclai as small (≤ 0.20), medium (>0.20 & < 0.60), large (≥ 0.60 & < 1.0) and largest (≥ 1.0). **B**. Distribution of all nuclei among packets (replotted from Fig. 4E). **C.** Distribution of small, medium, large and largest nuclei among packets; n indicates the total number of nuclei in each size class.

The distribution among packets of the smallest nuclei (! 0.20 on the normalized size scale) is about the same as for the distributions of neurons overall (cf. Fig. 7B and S3). In contrast, the three larger size classes show packet-specific distribution patterns (Fig. 7C); the medium size class (> 0.20 and < 0.60) is under-represented in AVM and maximally represented in PVM; the large size class (# 0.60 and < 1.0) is largely confined to the three anterior packets (LAL, AVM and RAL); and the largest nuclei (# 1.0) are mostly located in the LAL and RAL packets.

In comparing nuclei of different size classes, we note that the histone-labeled chromatin of some of the largest nuclei has the same or higher fluorescence intensity as in much smaller nuclei (data not shown). This observation is consistent with the hypothesis that different nuclear sizes reflect differences in cell ploidy, but differences in the mCherry signal could also reflect differences among cells in the level of *Hau-elav4*-dependent expression and/or differences in laser illumination during imaging. Thus, measuring differences in ploidy directly is beyond the scope of the present work.

## DISCUSSION

### Transposon-mediated transgenesis in leech

The implementation of genetic and molecular approaches has made it possible to carry out remarkably detailed investigations of intensively studied models such as mouse, zebrafish *Drosophila,* and *Caenorhabditis*. Inevitably, the very power and resource-intensity of these approaches have necessarily restricted the phylogenetic breadth of investigations in modern biology. In particular, these models represent just two of the three main branches of bilaterian animal diversity, the super-phyla Deuterostomia and Ecdysozoa.

The third main branch of animal evolution, the super-phylum Spiralia (Marlétaz et al., 2019), is of interest because it includes the plurality of modern animal phyla. This group is severely understudied compared to the other two, however, and the application of genome editing approaches is correspondingly sparse. Given their phylogenetic position, leeches provide a reference for studying how developmental processes have diverged among diverse animals over half a billion years of independent evolution.

At the time of this writing, stable transgenesis in Spiralia has been achieved by microinjecting zygotes for a mollusc, *Crepidula* (using CRISPR/Cas9 to achieve targeted knockins; Perry and Henry, 2015), for a polychaete annelid, *Platynereis* (using a Tc1/mariner-type transposable element; Backfisch et al., 2013), and for a planarian flatworm, *Macrostomum* (using circular or linear DNA fragments; Wudarski et al., 2022). Here, we add to the list of organisms in the superphylum Spiralia for which transgenic approaches have been applied successfully, using a Minos-based transposable element to efficiently introduce transgenes into the genome of a glossiphoniid leech, *Helobdella austinensis*. This is noteworthy because the Minos-based approach is quite efficient, and because *Helobdella* is already among the most tractable spiralian embryos. For example, whereas work in most other organisms is largely restricted to zygote injections, the *Helobdella* embryo is amenable to injecting identifiable, lineage-restricted stem cells or other cells to mark specific somatic lineages in the F0 animals, as well as generating stable transgenic lines by injecting germline precursors throughout early development.

A somewhat paradoxical aspect of our results is that while transgenesis is quite efficient, the copy number of insertions appears to be low. In a typical experiment, most of the surviving embryos have incorporated the transgene, and yet the inverse PCR analysis indicates that just one or two copies of the transgene were heritably inserted. One possible explanation for these results is that embryos with higher copy number insertions die during development.

Another possibility is that some intrinsic factors might restrict Minos-mediated insertions to certain “hotspots” in the *Helobdella* genome. At any rate, the fact that we can readily detect single-locus insertions of transgenes driven by the native *Hau-elav4* enhancer suggests that it should be feasible to detect reporters knocked into endogenous genes (expressed at comparable levels), for example by CRISPR/Cas9 mutagenesis coupled with homology-directed repair.

### Analysis of transgenic *Helobdella* reveals neuroanatomical compaction and increased variability compared to other leech species

Invertebrate species are used to investigate a wide range of neurobiological phenomena because they offer experimental advantages such as large cells (e.g., the squid giant axon and synapse) and/or cellular simplicity (e.g., *Caenorhabditis*). Invertebrates also provide fascinating examples of how nervous systems have diverged and specialized among diverse kinds of animals (Bullock and Horridge, 1965).

Individually identified neurons have been characterized in animals ranging in complexity from nematodes (e.g. *Caenorhabditis elegans*) to fish (Sillar, 2009). As an example of relevance here, the segmental ganglia of medicinal leeches (*Hirudo medicinalis* and *H. verbana*) have been used extensively to study neural circuitry and the properties of individual neurons (Calabrese et al., 2016; Szczupak, 2023). These studies rely in part on the ability to reproducibly identify individual neurons from ganglion-to-ganglion, animal-to-animal, and even among different leech species (Nusbaum and Kristan, 1986). Our work provides the groundwork for mapping identified neurons within the midbody segmental ganglia in *Helobdella*. Such maps will be useful for future neurobiological analyses in this species.

The extent to which such neural stereotypy exists for animals has not been extensively assessed, with few exceptions. For the nematode *Caenorhabditis elegans*, serial EM reconstruction revealed precisely 302 neurons in the hermaphrodite, in 118 morphological classes (White et al., 1986). The initial reconstruction was based on an overlapping combination of series from different regions of multiple individuals, but a large body of subsequent work has effectively confirmed those results. Earlier, serial reconstruction had been used to study the structure and projections of the cyclopic compound eye in an isogenic (parthenogenetic) strain of the water flea *Daphnia magna* (Macagno et al., 1973). These authors found an essentially invariant cellular structure to the eye, with constancy in the overall numbers of photoreceptors, in their organization into ommatidia and in the multiplicity of the projections of ommatidia to lamina cells in the optic lobe. Variability emerged in the details of branch morphologies and synaptic connections. Taken to the extreme, this previous work suggests that "simple" systems have rigidly fixed neural composition and largely stereotyped wiring of neural circuits.

For the leech nervous system, the existence of numerous individually identified neurons fits well with the deterministic view (Nicholls and Baylor, 1968). For example, a pair of large serotonergic neurons, known as Retzius (Rz) cells, are readily detected by their serotonin content throughout clitellates, i.e., oligochaetes and leeches (Koza et al., 2006); among leeches, the Rz neurons lie routinely in the AVM packet. Two other examples (from among many conserved cellular phenotypes) are the mechanosensory nociceptive (N) and pressure (P) neurons. Their cell bodies are about the same size, but N cells have their cell bodies in LAL and RAL packets, while the P cell bodies are located in LPL and RPL packets. Homologs of these characterized *Hirudo* neurons have been identified physiologically in other leech species including glossiphoniid species by both physiological and molecular approaches (Heath-Heckman et al., 2021; Kramer and Goldman, 1981).

A precise anatomical analysis revealed variation in the neuronal content of ganglia in several leech species, but only of a minor nature. Computer-assisted analyses of six serially sectioned and 20 whole mount ganglia from three hirudinid and one glossiphoniid leech species revealed that the neuronal content in ganglion M10 of *Hirudo medicinalis* ranged from 389 to 398 for an average of 394 ± 5 (Av ± SD) for and a coefficient of variation of 0.01 (Macagno, 1980). Two other hirudinid species, *Macrobdella decora* and *Haemopis marmorata*, were similar in range and consistency for ganglion M10. The same study found the same degree of consistency for *Haementeria ghilianii*, but this glossiphoniid species contained only around 380 neurons in ganglion M10 (Macagno, 1980).

In the present work, we have analyzed the numbers and sizes of neuronal nuclei (via the proxy of their nuclear chromatin) in segmental ganglia of a fifth leech species. *Helobdella austinensis* is a glossiphoniiid species like *Haementeria* only much smaller in body size. Our data suggest that the variability of neuronal architecture in *Helobdella* is far greater than found in four larger species. We find that *Helobdella* midbody segmental ganglia M02-M18 each contain about 320 neurons, significantly fewer than in the other four species. We also observe a greater range and variability in the number of neurons in *Helobdella* ganglia, vis-à-vis the other species. For example, our three counts of *Helobdella* ganglion M10 range from 310 to 348 with an average of 328 ± 19 neurons, for a coefficient of variation of 0.06, sixfold greater than that reported in *Hirudo* (Macagno, 1980). Other ganglia showed even greater ranges (Fig. 5C). No differences between the ganglia in genital segments (M5 and M6) and non-genital segments were observed for *Helobdella* in contrast to *Hirudo* and *Haementeria* where these segments contain significantly more neurons than the rest of the segments (Macagno 1980), but small differences could be masked by the large variability.

Here, we used transgenic approaches that are currently limited to *Helobdella*, using a pan-neuronal enhancer to drive a readily detected fluorescent protein marker, and the data were obtained by confocal microscopy. These methods allowed us to count a more complete sample of ganglia and eliminated the problem of distinguishing between neurons and non-neuronal cell types in the ganglia.

The differences in both methodology and species mean that we cannot unambiguously compare our results to previous work (Macagno, 1980). Thus, we cannot exclude the formal possibility that the observed variance arises artifactually from the imaging procedure or from genetic perturbations inherent to the transgenesis. These explanations seem unlikely, however. Regarding the former possibility, objects identified as neurons in some sample packets were validated by a second author, which argues against methodological errors, as does the high signal-to-noise ratio of the transgenic reporter. On the other hand, we have observed neither an increase in embryonic lethality for the transgenic line compared to wildtype animals (that would be evidence of developmental abnormalities), nor any behavior deficits in locomotion, feeding or reproduction (that would be evidence of malformation in the nervous system).

In addition to the differences discussed above, our more extensive data set confirms previous conclusions that the neuronal content of the six glia packets is also variable. Despite this variability, we observe conservation in the pattern of distribution of the large neurons. In addition to the pair of large serotonergic neurons (Retzius cells), that are almost always located in the AVM and which are apparently present throughout the Citellata (leeches and oligochaetes), *Helobdella* exhibits additional giant cells in the LAL and RAL packet of *Helobdella*. Based on the similarities between *Helobdella* and *Haementeria*, we assume that these giant cells represent a glossiphoniid-specific pattern, and that the >150-fold range of chromatin area/volume across *Helobdella* neurons represents a cell type specific rounds of post-mitotic DNA amplification (polyploidy). A question for future work will be to determine if distinct size classes correspond to particular cell types that undergo controlled amounts of post-mitotic DNA amplification as a part of their differentiation program.

### Evolutionary and developmental implications

One biologically based speculation explaining the observed differences (fewer neurons and higher variability) in *Helobdella* versus previously examined leech species is based on the dramatically smaller body size of *Helobdella* compared to *Haementeria* or *Hirudo*, and on the differences in scaling parameters for various biological functions. We speculate first that the smaller size means that it is not essential to have one pair of each type of neuron in each segment of the nerve cord; i.e., branches from one neuron of a given type in one segment might serve the functions of that same cell type in adjacent segments and/or on the contralateral side of the animal. Previous work has revealed such interactive and competitive interactions among segmentally homologous neurons in *Hirudo*, *Haementeria* and *Helobdella* (Blair et al., 1990; Gao and Macagno, 1987; Kramer et al., 1985; Kramer and Stent, 1985; Kuwada, 1985; Macagno and Stewart, 1987; Martindale and Shankland, 1990; Muller et al., 1987). If we further assume that "redundant" neurons die in a stochastic manner during normal neural maturation, it could account for both the reduced cell number and the increased variability in cell number between *Helobdella* and *Haementeria*. Assuming that this stochastic pruning of ganglionic cell numbers occurs only as the postmitotic neurons are extending their processes, a testable prediction of this model is that segmental ganglia in juvenile *Helobdella* could have larger and less variable numbers of cells, comparable to those in adult *Haementeria*.

Regardless of the underlying mechanism, the unexpectedly high variability we observed in the neuronal content of segmental ganglia in *Helobdella* suggests that this relatively simple nervous system is nonetheless capable of extensive neurodevelopmental plasticity. To investigate these issues with greater precision, it should be feasible to extend the experimental approach introduced here to examine specific subpopulations of neurons, using DNA elements capable of driving cell type-specific reporter expression. Thus, by complementing the rich knowledge of the leech nervous system based mainly on *Hirudo*, *Helobdella* is poised to become a promising model system to explore the nature of determinism and plasticity in a simple nervous system.

## Acknowledgements

The authors would like to thank Alexander D. Hartenstein (UC Berkeley) for his pilot work leading to the successful implementation of the Minos-based transgenic protocol in *Helobdella*, Wei-Yu Tao (NTU) for screening candidate genes for enhancer cloning, and Zhi-Yi Qiu (NTU) for his efforts in maintaining transgenic lines.

## Author contributions

Conceptualization: DHK, LS, DAW; Methodology: DHK, LS, DAW; Formal analysis: DHK, ELP, JRL, LS, FYT, CJW, DAW; Investigation: DHK, ELP, JRL, LS, FYT, CJW, DAW; Resources: DHK; Data curation: LS; Writing - original draft: DAW, DHK, LS; Writing - review & editing: CJW, DAW, DHK, ELP, LS; Visualization: CJW, DHK, ELP, LS; Funding acquisition: DHK, LS, DAW, ELP

## Data availability

Data, reagents and transgenic strains are available from the authors upon reasonable request.

## References

Backfisch, B., Rajan, V. B. V., Fischer, R. M., Lohs, C., Arboleda, E., Tessmar-Raible, K. and Raible, F. (2013). Stable transgenesis in the marine annelid *Platynereis dumerilii* sheds new light on photoreceptor evolution. Proc. Nat. Acad. Sci. U. S. A. 110, 193–198.

Blair, S. S., Martindale, M. Q. and Shankland, M. (1990). Interactions between adjacent ganglia bring about the bilaterally alternating differentiation of RAS and CAS neurons in the leech nerve cord. J. Neurosci. 10, 3183–3193.

Bullock, T. H. and Horridge, G. A. (1965). Structure and Function in the Nervous Systems of Invertebrates. San Francisco, CA: W. H. Freeman.

Calabrese, R. L., Norris, B. J. and Wenning, A. (2016). The neural control of heartbeat in invertebrates. Curr. Opin. Neurobiol. 41, 68–77.

Cho, S. J., Vallès, Y. and Weisblat, D. A. (2014). Differential expression of conserved germ line markers and delayed segregation of male and female primordial germ cells in a hermaphrodite, the leech *Helobdella*. Mol. Biol. Evol. 31, 341–354.

Coggeshall, R. E. and Fawcett, D. W. (1964). The fine structure of the central nervous system of the leech, *Hirudo medicinalis*. J. Neurophysiol. 27, 229–289.

Colombrita, C., Silani, V. and Ratti, A. (2013). ELAV proteins along evolution: back to the nucleus? Mol. Cell Neurosci. 56, 447–455.

Edgar, R. C. (2022). Muscle5: high-accuracy alignment ensembles enable unbiased assessments of sequence homology and phylogeny. Nat. Commun. 13, 6968.

Gao, W.-Q. and Macagno, E. R. (1987). Extension and retraction of axonal projections by some developing neurons in the leech depends upon the existence of neighboring homologues. II. The AP and AE neurons. J. Neurobiol. 18, 295–313.

Gline, S. E., Kuo, D.-H., Stolfi, A. and Weisblat, D. A. (2009). High resolution cell lineage tracing reveals developmental variability in leech. Dev. Dyn. 238, 3139–3151.

Heath-Heckman, E., Yoo, S., Winchell, C., Pellegrino, M., Angstadt, J., Lammardo, V. B., Bautista, D., De-Miguel, F. F. and Weisblat, D. (2021). Transcriptional profiling of identified neurons in leech. BMC Genomics 22, 215.

Hoang, D. T., Chernomor, O., von Haeseler, A., Minh, B. Q. and Vinh, L. S. (2018). UFBoot2: improving the ultrafast bootstrap approximation. Mol. Biol. Evol. 35, 518–522.

Kalyaanamoorthy, S., Minh, B. Q., Wong, T. K. F., von Haeseler, A. and Jermiin, L. S. (2017). ModelFinder: fast model selection for accurate phylogenetic estimates. Nat. Methods 14, 587–589.

Kang, D., Pilon, M. and Weisblat, D. A. (2002). Maternal and zygotic expression of a *nanos*-class gene in the leech *Helobdella robusta*: primordial germ cells arise from segmental mesoderm. Dev. Biol. 245, 28–41.

Koza, A., Wilhelm, M., Hiripi, L., Elekes, K. and Csoknya, M. (2006). Embryogenesis of the serotonergic system in the earthworm Eisenia fetida (Annelida, Oligochaeta): immunohistochemical and biochemical studies. J. Comp. Neurol. 497, 451–467.

Kramer, A. P. and Goldman, J. R. (1981). The nervous system of the glossiphoniid leech *Haementeria ghilianii*. I. Identification of neurons. J. Comp. Physiol. A 144, 435–448.

Kramer, A. P., Goldman, J. R. and Stent, G. S. (1985). Developmental arborization of sensory neurons in the leech *Haementeria ghilianii*. I. Origin of natural variations in the branching pattern. J. Neurosci. 5, 759–767.

Kramer, A. P. and Stent, G. S. (1985). Developmental arborization of sensory neurons in the leech *Haementeria ghilianii*. II. Experimentally induced variations in the branching pattern. J. Neurosci. 5, 768–775.

Kramer, A. P. and Weisblat, D. A. (1985). Developmental neural kinship groups in the leech. J. Neurosci. 5, 388–407.

Kuo, D.-H., De-Miguel, F. F., Heath-Heckman, E. A. C., Szczupak, L., Todd, K., Weisblat, D. A. and Winchell, C. J. (2020). A tale of two leeches: toward the understanding of the evolution and development of behavioral neural circuits. Evol. Dev. 22, 471–493.

Kuo, D.-H. and Weisblat, D. A. (2011). Intermediate filament genes as differentiation markers in the leech *Helobdella*. Dev. Genes Evol. 221, 225–240.

Kuwada, J. Y. (1985). Pioneering and pathfinding by an identified neuron in the embryonic leech. J. Embryol. Exp. Morphol. 86, 155–167.

Lasek, R. J. and Dower, W. J. (1971). *Aplysia californica*: analysis of nuclear DNA in individual nuclei of giant neurons. Science 172, 278–280.

Macagno, E. R. (1980). Number and distribution of neurons in leech segmental ganglia. J. Comp. Neurol. 190, 283–302.

Macagno, E. R., Lopresti, V. and Levinthal, C. (1973). Structure and development of neuronal connections in isogenic organisms: variations and similarities in the optic system of *Daphnia magna*. Proc. Nat. Acad. Sci. U. S. A. 70, 57–61.

Macagno, E. R. and Stewart, R. R. (1987). Cell death during gangliogenesis in the leech: competition leading to the death of PMS neurons has both random and nonrandom components. J. Neurosci. 7, 1911–1918.

Marlétaz, F., Peijnenburg, K. T. C. A., Goto, T., Satoh, N. and Rokhsar, D. S. (2019). A new spiralian phylogeny places the enigmatic arrow worms among gnathiferans. Curr. Biol. 29, 312–318.

Martindale, M. Q. and Shankland, M. (1990). Neuronal competition determines the spatial pattern of neuropeptide expression by identified neurons of the leech. Dev. Biol. 139, 210–226.

McGlade-McCulloh, E., Morrissey, A. M., Norona, F. and Muller, K. J. (1989). Individual microglia move rapidly and directly to nerve lesions in the leech central nervous system. Proc. Nat. Acad. Sci. U. S. A. 86, 1093–1097.

Muller, K. J., McGlade-McCulloh, E. and Mason, A. (1987). Tinkering with successful synapse regeneration in the leech: adding insult to injury. J. Exp. Biol. 132, 207–221.

Muller, K. J., Nicholls, J. G. and Stent, G. S. (1981). Neurobiology of the Leech, pp. 320. Cold Spring Harbor, NY: Cold Spring Harbor Laboratory Press.

Nguyen, L.-T., Schmidt, H. A., von Haeseler, A. and Minh, B. Q. (2015). IQ-TREE: a fast and effective stochastic algorithm for estimating maximum-likelihood phylogenies. Mol. Biol. Evol. 32, 268–274.

Nicholls, J. G. and Baylor, D. A. (1968). Specific modalities and receptive fields of sensory neurons in CNS of the leech. J. Neurophysiol. 31, 740–756.

Nusbaum, M. P. and Kristan, W. B., Jr. (1986). Swim initiation in the leech by serotonin-containing interneurones, cells 21 and 61. J. Exp. Biol. 122, 277–302.

Pascale, A., Amadio, M. and Quattrone, A. (2008). Defining a neuron: neuronal ELAV proteins. Cell. Mol. Life Sci. 65, 128–140.

Pavlopoulos, A. and Averof, M. (2005). Establishing genetic transformation for comparative developmental studies in the crustacean *Parhyale hawaiensis*. Proc. Nat. Acad. Sci. U. S. A. 102, 7888–7893.

Pavlopoulos, A., Oehler, S., Kapetanaki, M. G. and Savakis, C. (2007). The DNA transposon *Minos* as a tool for transgenesis and functional genomic analysis in vertebrates and invertebrates. Genome Biol. 8, S2.

Pende, M., Vadiwala, K., Schmidbaur, H., Stockinger, A. W., Murawala, P., Saghafi, S., Dekens, M. P. S., Becker, K., Revilla-i-Domingo, R., Papadopoulos, S.-C. et al. (2020). A versatile depigmentation, clearing, and labeling method for exploring nervous system diversity. Sci. Adv. 6, eaba0365.

Perry, K. J. and Henry, J. Q. (2015). CRISPR/Cas9-mediated genome modification in the mollusc, *Crepidula fornicata*. genesis 53, 237–244.

Portiansky, E. L. (2018). Análisis multidimensional de imágenes digitales. La Plata: Universidad Nacional de La Plata. http://sedici.unlp.edu.ar/handle/10915/70938

Retzius, G. (1891). Zur Kenntnis des zentralen Nervensystems der Würmer. Biol. Unters. (N.F.) 2, 1–28.

Retzius, G. (1898). Zur Kenntniss des sensiblen Nervensystems der Hirudineen. Biol. Unters. (N.F.) 8, 94–97.

Shankland, M. and Savage, R. M. (1997). Annelids, the segmented worms. In Embryology: Constructing the Organism, eds. S. F. Gilbert and A. M. Raunio), pp. 219–235. Sunderland, MA: Sinauer.

Sillar, K. T. (2009). Mauthner cells. Curr. Biol. 19, R353–R355.

Szczupak, L. (2023). Motor neural networks in the leech. Trends Neurosci. 46, 698–700.

Wagenaar, D. A. (2015). A classic model animal in the 21st century: recent lessons from the leech nervous system. J. Exp. Biol. 218, 3353–3359.

Weisblat, D. A. (2022). Glossiphoniid leeches as a touchstone for studies of development in clitellate annelids. Curr. Top. Dev. Biol. 147, 433–468.

Weisblat, D. A., Kim, S. Y. and Stent, G. S. (1984). Embryonic origins of cells in the leech *Helobdella triserialis*. Dev. Biol. 104, 65–85.

Weisblat, D. A. and Kuo, D.-H. (2009). *Helobdella* (Leech): a model for developmental studies. In Emerging Model Organisms: a Laboratory Manual, pp. 245–267. Cold Spring Harbor, NY: CSHL Press.

Weisblat, D. A. and Kuo, D.-H. (2014). Developmental biology of the leech *Helobdella*. Int. J. Dev. Biol. 58, 429–443.

White, J. G., Southgate, E., Thomson, J. N. and Brenner, S. (1986). The structure of the nervous system of the nematode *Caenorhabditis elegans*. Philos. Trans. R. Soc. Lond. B. 314, 1–340.

Whitman, C. O. (1878). The embryology of *Clepsine*. Q. J. Microsc. Sci. 18, 213–315.

Whitman, C. O. (1887). A contribution to the history of the germ-layer in *Clepsine*. J. Morphol. 1, 105–182.

Wudarski, J., Simanov, D., Ustyantsev, K., de Mulder, K., Grelling, M., Grudniewska, M., Beltman, F., Glazenburg, L., Demircan, T., Wunderer, J. et al. (2017). Efficient transgenesis and annotated genome sequence of the regenerative flatworm model *Macrostomum lignano*. Nat. Commun. 8, 2120.

Yamagishi, M., Ito, E. and Matsuo, R. (2011). DNA endoreplication in the brain neurons during body growth of an adult slug. J. Neurosci. 31, 5596–5604.

Zhang, S. O. and Weisblat, D. A. (2005). Applications of mRNA injections for analyzing cell lineage and asymmetric cell divisions during segmentation in the leech *Helobdella robusta*. Development 132, 2103–2113..

